# Unraveling the Toxicological Effects of Hydroxyacetone - A Reaction Product in Electronic Cigarette Aerosols

**DOI:** 10.1101/2025.08.29.673133

**Authors:** Man Wong, Teresa Martinez, My Hua, Nathan Hendricks, Prue Talbot

## Abstract

Hydroxyacetone has been detected at high concentrations (up to ∼12 mg/mL) in electronic cigarette (EC) aerosols, including those derived from products associated with adverse health effects. Given the limited understanding of its inhalation toxicology, we investigated hydroxyacetone’s impact on human airway epithelial cells. Acute exposures at the air–liquid interface (ALI) using 3D EpiAirway tissues—a surrogate for human tracheobronchial epithelium—were analyzed via proteomics. Differential expression analysis identified numerous affected proteins, with enrichment pointing to alterations in mitochondrial function and actin cytoskeletal disruption as major targets. Ingenuity Pathway Analysis (IPA) highlighted “Mitochondrial Dysfunction” and “NRF2-Mediated Oxidative Stress” among top toxicological categories, and “Nuclear Cytoskeletal Signaling” as a key canonical pathway. To validate and extend these findings, submerged cultures of BEAS-2B cells were exposed to hydroxyacetone (0.01–10 mg/mL) and assessed for mitochondrial activity, oxidative stress, and F-actin integrity. At 1 mg/mL, mitochondrial membrane potential and reactive oxygen species (ROS) increased, with elevated hydrogen peroxide detected in the culture medium. At 10 mg/mL, mitochondrial activity declined significantly, accompanied by cell rounding and apoptotic blebbing within 2 hours. F-actin destabilization occurred at 1, 3.33, and 10 mg/mL, with cytoplasmic and perinuclear filaments more affected than cortical actin. Findings from ALI and submerged models were concordant, supporting hydroxyacetone-induced mitochondrial stress, oxidative damage, and cytoskeletal disruption. These results suggest that hydroxyacetone concentrations found in EC aerosols may contribute to respiratory toxicity and warrant further investigation.

## INTRODUCTION

Electronic-cigarette (EC) use is associated with a wide range of self-reported symptoms, such as nausea, headache, heart palpitations, dizziness, chest pain, and cough, that affect the respiratory, gastrointestinal, neurological and immune systems (1,2). More serious conditions, often physician-diagnosed (signs) and sometimes based on case reports, have also been described, including seizures, myocardial infarction, bronchitis, dermatitis, burns, and nicotine poisoning (3–5). Much of this literature has appeared in systematic and umbrella reviews (6–9). In 2020, ECs were linked to the severe respiratory disease E-Cigarette Vaping Associated Lung Injury (EVALI) epidemic (10). While many cases of EVALI were associated with product contamination by vitamin E acetate (11,12), the etiology of a subset of cases remains unknown (13) and underreporting of EVAI cases is a concern (14).

Following the EVALI epidemic, reports continued to document adverse health effects associated with EC use (15,16), yet the causes of EC-induced illness remain incompletely understood. A meta-analysis identified health outcomes in EC users that were distinct from those observed in cigarette smokers (16), and a recent umbrella review summarized EC-related health issues reported in adolescents and young adults (17). These findings underscore the need to investigate the chemical constituents of EC aerosols and their biological effects.

Most EC liquids contain varying concentrations of nicotine, flavor chemicals, and solvents, such as propylene glycol and glycerin, which are aerosolized and inhaled during vaping (18–21). Fourth-generation ECs and newly introduced ultrasonic SURGE devices also contain high levels of synthetic coolants, including WS-23 (22–25), which impairs cytoskeletal function in human respiratory epithelium (26). Common flavor chemicals include vanillin, ethyl maltol, ethyl vanillin, and menthol (27,28). Although many of these compounds are generally recognized as safe (GRAS) for ingestion, the Flavor and Extract Manufacturers Association (FEMA) has not evaluated their safety for inhalation, and their toxicological profiles when aerosolized remain poorly characterized (29). In vitro and in vivo studies suggest that several flavor chemicals, including maltol, ethyl maltol, vanillin, cinnamaldehyde, and menthol, can be harmful when inhaled at the high concentrations found in EC products (27, 30–35).

EC aerosols also contain benzoic acid, ultrafine particles, metals, and numerous reaction products formed during storage or heating of flavor chemicals, propylene glycol, and glycerin (36–43). Many of these reaction products are toxic and appear on the FDA’s list of Harmful and Potentially Harmful Constituents (HPHC) (44). Unlike flavor chemicals added at known concentrations, reaction products formed during aerosolization are often unidentified, unknown to the user, and highly variable across users and devices (39,45). Some, such as formaldehyde are known carcinogens (46), and others, like diacetyl, have been linked to epithelial injury and bronchiolitis obliterans (47).

The present study investigates the toxicological effects of a specific reaction product, hydroxyacetone, on human airway epithelial cells. Hydroxyacetone was selected based on its prevalence and abundance in EC aerosols generated from 140 refill fluid products (48). It was detected in nearly all (126) aerosol samples and reached concentrations as high as 12 mg/mL - orders of magnitude higher than other reaction products identified in the same study. Despite its widespread presence in EC aerosols, little is known about the toxicity of hydroxyacetone in the context of inhalation exposure.

To address this gap, we first conducted proteomic analysis using 3D EpiAirway tissues exposed to hydroxyacetone at the air–liquid interface (ALI) to obtain a global overview of cellular responses and identify targets. EpiAirway is a differentiated human tracheobronchial tissue grown in vitro from airway basal stem cells. It replicates the structure and function of the native airway epithelium and recapitulates the *in vivo* phenotypes of barrier, mucociliary responses, infection, toxicity responses, and disease. Based on proteomic findings, follow-up experiments were performed using BEAS-2B cells in monolayer culture to validate key targets and further characterize cellular responses to hydroxyacetone. Together, these studies provide new insight into the biological impact of hydroxyacetone on airway epithelial integrity and function.

## METHODS

### EC Products and Acquisition

A sample set of 140 EC refill fluids was purchased from online and local vape shops based on previously reported data (48). These EC fluids had been reported online by users to produce numerous symptoms including, but not limited to, nausea, burning throat, chest pain, irritated throat, cough, lightheadedness, headache, and wheezing. Samples were stored at 4°C until they were used to make aerosols. A complete list of products included in this sample set is provided in Supplementary Table 1.

Prior chemical analysis of aerosols generated from these fluids identified five reaction products. Notably, hydroxyacetone was consistently detected in the aerosols despite being absent from the unvaped fluids, with concentrations reaching up to 12 mg/mL. In contrast, diacetyl, acetophenone, furfural, and strawberry glycidate were detected in a limited subset of aerosols, typically at concentrations below 0.5 mg/mL (48). Given its consistent presence and limited prior investigation in the context of inhalation toxicology, hydroxyacetone was selected for further study.

### VITROCELL® Analysis Cloud Chamber Exposures of EpiAirway Tissues for Proteomics

Exposure of EpiAirway™ tissues to hydroxyacetone aerosols was done in a VITROCELL® cloud chamber (VITROCELL® 12/12 base module, Walkirch, Germany). EpiAirway maintenance medium was equilibrated to 37°C in each well of the cloud chamber for 15 minutes before exposure. Aerosols were generated using an Aerogen vibrating mesh nebulizer (AG-AL1100, San Mateo, California). 200 µL of either PBS- (phosphate buffered saline minus calcium and magnesium) or 10 mg/mL of hydroxyacetone (138185, Sigma-Aldrich) dissolved in PBS-, were loaded into the nebulizer to generate an aerosol with minimal thermal degradation of the solutions.

Tissues were exposed to one puff of either PBS-control aerosol or hydroxyacetone aerosol, returned to the incubator for 4 hours, then exposed to a second puff, after which tissues were returned to the incubator for a 24-hour recovery period prior to proteomic analysis. This exposure protocol was designed to align with prior studies conducted in our laboratory (26, 34) and to approximate a realistic vaping exposure scenario. The 24-hour post-exposure interval was selected to allow for protein expression changes in response to chemical treatment.

### Proteomics Sample Preparation

Lysed EpiAirway tissues were adjusted to 500 µL with 50 mM triethylammonium bicarbonate (TEAB, Sigma Aldrich, St. Louis, MO). Proteins were precipitated by adding an equal volume of trichloroacetic acid and incubated on ice for 1 hour. The resulting supernatant was discarded, and the protein pellets were washed twice with 500 µL of cold acetone, which were also discarded. Residual acetone was removed by vacuum centrifugation (SpeedVac).

Dried pellets were resuspended in 50 µL of in 50 mM TEAB, then reduced with 1 μL of 500 mM tris(2-carboxyethyl)phosphine (Thermo Scientific, Rockford, IL) and incubated at 37°C for 1 hour. 3 µL of 500 mM Iodoacetamide (Sigma Aldrich) were added to each sample, which were incubated in the dark at room temperature for 1 hour. 500 µL of 50 mM TEAB were then added to the samples, along with trypsin/lysC mix (Promega, Madison, WI) at a 1:80 ratio to protein mass. Samples were digested at 37°C overnight (16 hours). The resulting peptide solution was cleaned with Waters HLB solid-phase extraction columns.

Peptide concentrations were quantified using a colorimetric peptide assay (Thermo Scientific), and samples were grouped into a TMT 10-plex runs. For labeling, 3 µg of peptide from each sample was aliquoted and adjusted to pH 8 using 1 M TEAB. TMT reagents (Thermo Scientific) were added at a 1:1 mass ratio and incubated at room temperature for 1 hour. Labeling reactions were quenched with 8 µL of 5% hydroxylamine for 15 minutes.

Labeled samples were pooled and dried by SpeedVac prior to high-pH fractionation. The combined sample was resuspended in 0.1% trifluoroacetic acid and fractionated into eight fractions using the Pierce High-pH Reversed-Phase Peptide Fractionation Kit (Thermo Scientific). Each fraction was dried by SpeedVac and resuspended in 20 µL of 0.1% formic acid for liquid chromatography–mass spectrometry (LC-MS) analysis.

### Proteomics Analysis

Liquid chromatography was performed on a Thermo nLC1200 system in single-pump trapping mode, equipped with a PepMap RSLC C18 EASY-spray analytical column (2 μm, 100 Å, 75 μm x 25 cm) and with a PepMap C18 trap column (3 μm, 100 Å, 75 μm x 20 mm). Mobile phases were solvent A (water with 0.1% formic acid) and solvent B (80% acetonitrile with 0.1% formic acid). Peptide separation was conducted at a flow rate of 300LnL/min using a 130-minute gradient: 3% to 30% B from 1 to 110 minutes, ramping to 85% B at 120 minutes, followed by a 10-minute hold.

Mass spectrometry data were acquired on a Thermo Scientific Orbitrap Fusion instrument operating in data-dependent acquisition mode. Full scans were performed in the Orbitrap using 60,000 resolution in the positive mode. Precursors for MS2 were selected based on monoisotopic peak determination, with an intensity threshold of 5.0L×L10³, charge states of 2–7, and a dynamic exclusion window of 60 seconds following a single analysis, using a mass tolerance of 10Lppm. MS2 spectra were acquired in the ion trap using collision-induced dissociation (CID) at 35% normalized collision energy and an isolation window of 1.6Lm/z. Tandem Mass Tag (TMT) quantification was performed using synchronous precursor selection (SPS)-MS3 with high-energy collision dissociation (HCD) at 65% energy.

Raw data were processed using Proteome Discoverer 2.2 (Thermo Scientific) and searched against the UniProt FASTA database for Homo sapiens. Search parameters included a precursor mass tolerance of 10Lppm and fragment mass tolerance of 0.6LDa. Fixed modifications included carbamidomethylation of cysteine (+57.021LDa) and TMT labeling of lysine residues and peptide N-termini (+229.163LDa). Dynamic modifications included methionine oxidation (+15.995LDa) and N-terminal acetylation (+42.011LDa). Peptide-spectrum matches were filtered to a strict false discovery rate (FDR) of 1%.

Reporter ion abundances from MS3 spectra were normalized to total peptide abundance and scaled relative to the control sample across runs. Protein-level ratios were calculated, and statistical significance was assessed using one-way ANOVA across biological replicates. Proteomics excel files with absorbance values, Log2FC, p-values, and adjusted p-values are attached as an excel file (supplementary table 2).

### Gene Ontology (GO) Analysis

Protein lists for control and hydroxyacetone exposed groups were uploaded to the Gene Ontology (GO) database for functional annotation. Statistically significant proteins were identified using an adjusted p-value threshold of < 0.05. GO terms from the three primary ontologies (Biological Process, Cellular Compartment, and Molecular Function) were extracted and grouped using the rrvgo package in R (49). This package clusters semantically similar GO terms and designates a representative “parent term” based on statistical significance. Treemap visualizations of grouped GO terms were generated using the ggplot2 package in R.

### Ingenuity Pathways Analysis (IPA)

The Ingenuity Pathway Analysis (IPA) software was used to further investigate the biological processes and pathways affected by hydroxyacetone exposures at the ALI. The IPA “ToxList” tool used to associate experimental data to clinical pathology endpoints relevant to xenobiotic responses. Canonical Pathway analysis was also performed to identify signaling networks potentially impacted by hydroxyacetone. Circle plots mapping proteins to IPA pathways, ToxList functions, and GO terms were generated in R using the circlize package.

### Quantifying Hydroxyacetone Exposures in ALI Transwell Inserts

To quantify the concentration of hydroxyacetone deposited onto transwell inserts during aerosol exposure in the VITROCELL® cloud chamber, a single puff of aerosol was directed onto 1 mL of isopropyl alcohol placed in each insert. The resulting solutions were aliquoted into GC/MS vials and submitted to Portland State University for chemical analysis. Hydroxyacetone concentrations were determined via gas chromatography–mass spectrometry (GC/MS), and mean values with standard deviations were calculated for each transwell sample (Supplementary Figure 1A).

### BEAS-2B Cell Culture and Hydroxyacetone Exposure

BEAS-2B human bronchial epithelial cells were cultured in serum-free Bronchial Epithelial Growth Medium (BEGM; Lonza), supplemented with growth factors as previously described (27). Culture flasks (Corning Inc., Corning, NY) were pre-coated with Bronchial Epithelial Basal Medium (BEBM) containing collagen (30 µg/mL), fibronectin (10 µg/mL), and bovine serum albumin (BSA; 10 µg/mL). Cells were maintained at 37 °C in a humidified incubator with 5% CO₂, at 30–90% confluence. Subculturing was performed every 2–4 days using trypsin-mediated dissociation.

### MTT Assay

Mitochondrial activity was assessed using the methyl thiazolyl tetrazolium (MTT) assay. BEAS-2B cells (10,000 cells/well) were plated in 96-well plates and exposed to hydroxyacetone for 24 hours under submerged culture conditions. Following treatment, the culture medium was removed and replaced with MTT solution, followed by a 2-hour incubation at 37 °C. The MTT solution was then aspirated, and 200 µL of dimethyl sulfoxide (DMSO) was added to each well to solubilize the formazan crystals. Absorbance was measured at 570 nm using a Synergy HTX Microplate Reader (BioTek, Winooski, VT). Mean absorbance values and standard errors of the mean (SEM) were used to generate concentration–response curves.

### Mitochondrial Membrane Potential Assessment Using MitoTracker™ Deep Red

Mitochondrial activity was assessed using MitoTracker™ Deep Red FM (Thermo Fisher Scientific, Cat. No. M22426), a fluorescent dye that selectively accumulates in mitochondria in proportion to membrane potential. BEAS-2B cells were seeded into 6-well IBIDI µ-Slide chambers and exposed to hydroxyacetone at concentrations of 0.01, 0.1, 1, and 10 mg/mL for 24 hours. Following treatment, cells were incubated with MitoTracker™ Deep Red according to the manufacturer’s protocol. Live-cell imaging was performed using a Nikon Eclipse inverted fluorescence microscope to visualize mitochondrial fluorescence intensity.

### Mitochondrial Superoxide Detection Using MitoSOX™ Red

Mitochondrial superoxide production was assessed using MitoSOX™ Red (Thermo Fisher Scientific, Cat. No. M36007), a fluorogenic dye that selectively targets mitochondria and fluoresces upon oxidation by superoxide. BEAS-2B cells were seeded into 8-well IBIDI µ-Slide chambers and treated with hydroxyacetone at concentrations of 0.01, 0.1, 1, and 10 mg/mL for 24 hours. Following exposure, cells were incubated with MitoSOX™ Red according to the manufacturer’s protocol. Live-cell fluorescence imaging was performed using a Nikon Eclipse inverted fluorescence microscope.

### Quantification of Hydrogen Peroxide in Culture Medium Using the Amplex® Red Hydrogen Peroxidase Assay

Hydrogen peroxide (H₂O₂) levels in culture medium were quantified using the Amplex® Red Hydrogen Peroxidase Assay Kit (Thermo Fisher Scientific), a fluorometric assay based on the 1:1 stoichiometric reaction between Amplex® Red reagent and H₂O₂ in the presence of horseradish peroxidase (HRP), yielding the fluorescent product resorufin. BEAS-2B cells were exposed to hydroxyacetone at concentrations of 0.01, 0.1, and 1Lmg/mL for 24 hours. A standard curve was generated using serial dilutions of H₂O₂ (0–5 µM) in the kit’s reaction buffer, pipetted in duplicate into a black 96-well fluorescence plate. Fresh culture medium samples were loaded undiluted into designated wells, followed by the addition of Amplex® Red reagent and HRP to each well (excluding blanks). Plates were incubated at room temperature for 30 minutes in the dark. Fluorescence intensity was measured at an excitation/emission wavelength of 490/590 nm using a Synergy microplate reader (Agilent Technologies), and H₂O₂ concentrations were interpolated from the standard curve.

### Assessment of F-Actin Organization Following Hydroxyacetone Exposure Using Phalloidin-iFluor 594

BEAS-2B cells were seeded into Ibidi™ µ-Slide chamber slides and incubated at 37 °C in a humidified atmosphere containing 5% CO₂ for 24 hours. Cells were then treated with hydroxyacetone at concentrations of 1, 3.3, or 10Lmg/mL for 8 hours. Following treatment, cells were rinsed three times with phosphate-buffered saline (PBS), fixed with 4% paraformaldehyde for 10 minutes, and rinsed twice with PBS. F-actin filaments were labeled using Phalloidin-iFluor 594 (Abcam, Cat. No. ab176757) at a 1× working concentration, following the manufacturer’s protocol. Fluorescence imaging was performed using a Nikon Eclipse inverted fluorescence microscope equipped with a 60× objective and an Andor Zyla VSC-04941 camera. Images were acquired under non-saturating conditions and subsequently deconvoluted using Nikon Elements software.

### Statistical Analysis

For MTT and hydrogen peroxide data collected with BEAS-2B cells, means were compared using a one-way analysis of variance (ANVOA). When significance was detected, treated groups were compared to the untreated control using Dunnett’s post hoc test. If data did not meet the assumptions of ANOVA (homogeneity of variances and normal distribution), they were transformed using log(y) and rerun. Statistics were done using GraphPad Prism.

## RESULTS

### Proteomics of Hydroxyacetone Treated EpiAirway™ Tissues

Proteins with an adjusted p-value < 0.05 and a log2 fold-change > 0.585 or < -0.585 were considered significantly differentially expressed proteins (DEPs). A total of 107 downregulated DEPs and 193 upregulated DEPs were identified in the proteomics analysis (Figure 1A) and were included in the GO and IPA analyses. The top 20 DEPs, which are graphed in Supplementary Figure 1B, show that a broad range of processes (e.g., metabolism and biogenesis of lysosome-related organelles) were affected by exposure to hydroxyacetone.

**Figure 1:**
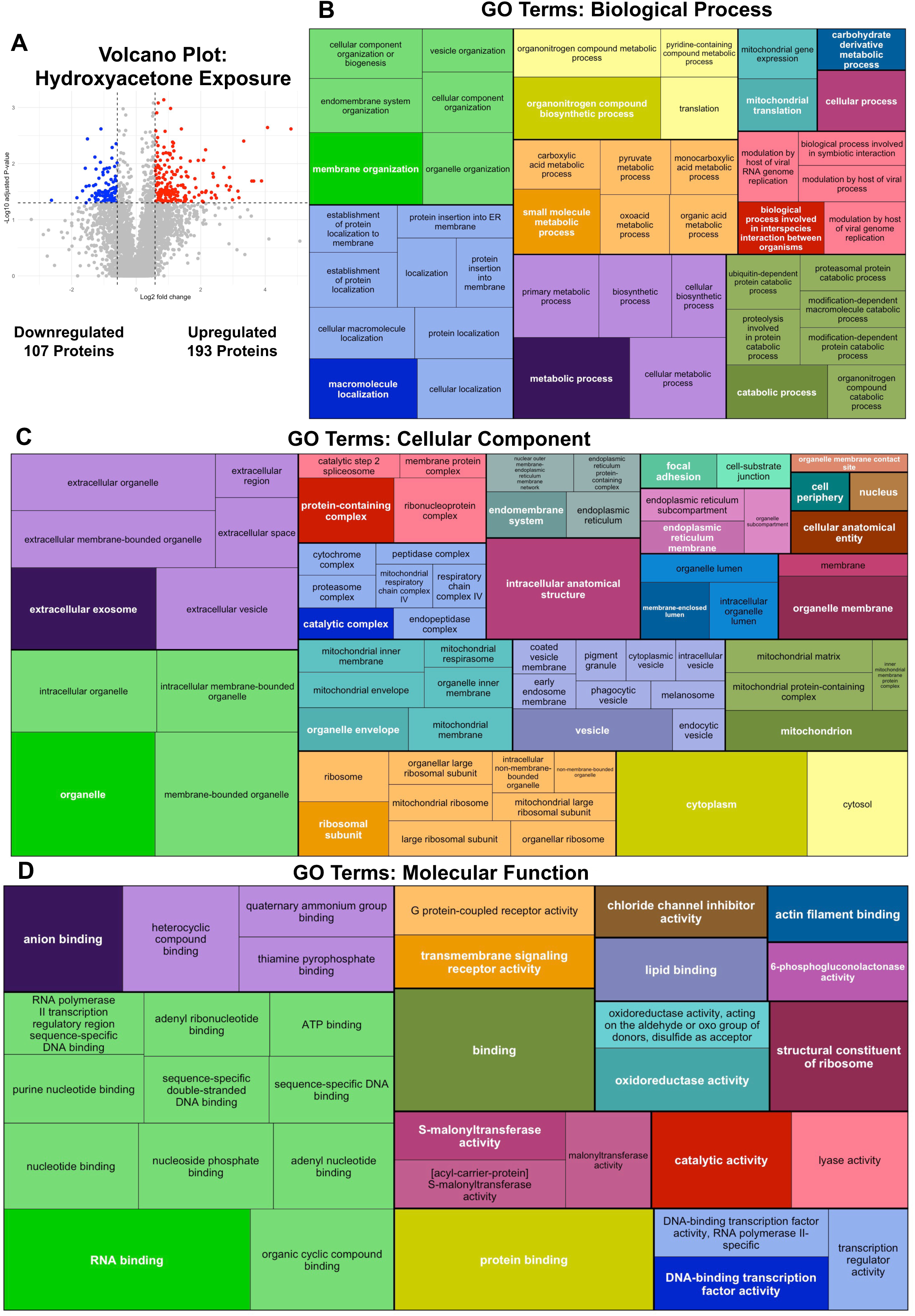
Volcano plot and GO terms for hydroxyacetone treated EpiAirway. DEPs in proteomics data were visualized in volcano plots (A) and treemaps (B-D) of the GO terms. In the volcano plot, red and blue points show up and downregulated DEPs, respectively. Treemap plots show GO terms for (B) Biological Process, (C) Cellular Compartment, and (D) Molecular Function. GO terms were grouped based on similarity using the rrvgo package. Each group has one parent GO term that is bolded and darker in color than other terms in the group. Within individual treemaps, box size corresponds to significance (–log10(p-value)) with more significant terms having larger sized boxes.

### GO Terms From Hydroxyacetone Treatments

To further characterize the effects of hydroxyacetone, GO analysis of DEPs was performed. GO Terms for the ontologies Biological Process, Cellular Component, and Molecular Function were plotted with the R package rrvgo (49), which groups similar GO terms together and designates the term with the highest significance (lowest adjusted p-value) as the parent term.

In the Biological Process GO Terms, the main effects involved in general metabolic related processes and energy production and included terms related to cell membrane organization and macromolecule localization, metabolic processes, and mitochondria (Figure 1B). The Cellular Component GO Terms indicated that the DEPs in our hydroxyacetone treatments were localized in many organelle compartments, including exosomes, organelle envelops, ribosomes, vesicles, and the mitochondria and mitochondrial catalytic complex (Figure 1C). GO Terms from the Molecular function ontology indicated that RNA binding was a major function effected by hydroxyacetone, in addition to anion binding, transmembrane signaling, DNA-binding transcription factor activity, and oxidoreductase activity (Figure 1D).

### IPA Analysis of Hydroxyacetone Treated EpiAirway™

IPA Tox List Analysis identified six significant Tox List functions from hydroxyacetone treated EpiAirway: Mitochondrial Dysfunction, Hypoxia-Inducible Factor Signaling, Aryl Hydrocarbon Receptor Signaling, NRF-2 Mediated Oxidative Stress Response, Glutathione Depletion-Phase II Reactions, and Increases Transmembrane Potential of Mitochondrial Membrane. A circle plot was generated connecting the Tox List functions to their corresponding DEPs (Figure 2A). The majority of the Tox List results identified mitochondria and oxidative stress as major targets of hydroxyacetone.

**Figure 2:**
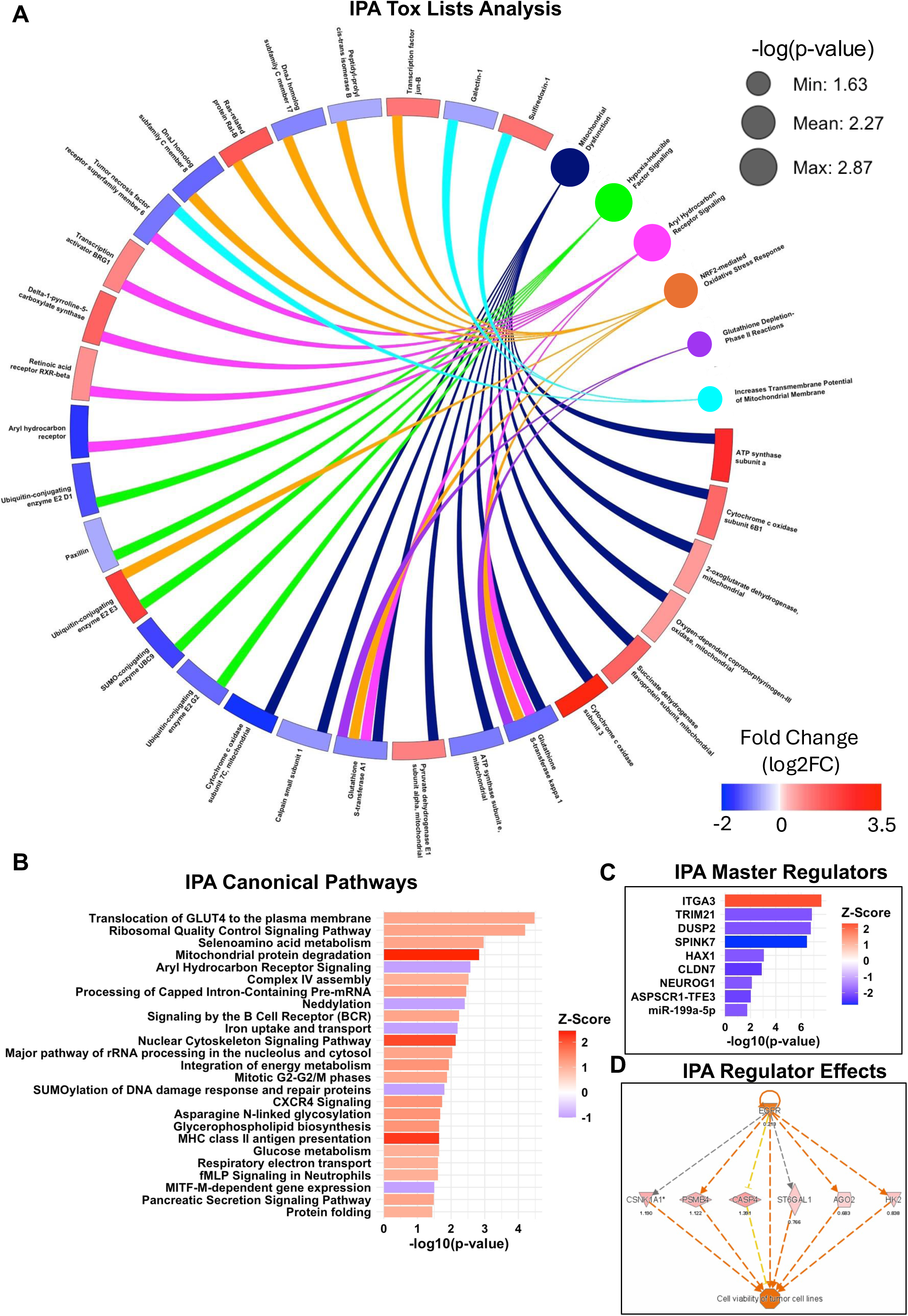
IPA Enrichment Analysis of hydroxyacetone treatments. (A) Circle plot connecting DEPs to IPA Tox List analysis. The size of the dots corresponding to Tox List function represents the -log10(p-value) of the pathway. The colored scale bar indicates the magnitude of the fold change for different proteins. (B) IPA Canonical Pathway Analysis with a z-score filter of +/-1. (C) Predicted IPA Master Regulators upstream of DEPs in hydroxyacetone treatments. (D) IPA Regulator Effects indicating EGFR as an upstream regulator of some DEPS.

IPA identified 25 canonical pathways (z > |1|) related to Ribosomal Signaling, Mitochondria, Cellular Respiration, Nuclear Cytoskeleton, Cell cycle, Immune Pathways, and Protein Folding (Figure 2B). The top three canonical pathways (z > 2) included Mitochondrial Protein Degradation, Nuclear Cytoskeleton Signaling Pathway, and MHC Class II Antigen Presentation; and are plotted in a circle plot with the pathways connecting to the DEPs (Supplemental Figure 2).

IPA Master Regulator analysis identified proteins that act upstream of the regulators of the DEPs identified in our dataset. Among these were the transcription factor TRIM21, which controls immune responses, and the mitochondrial protein HAX1, which interacts with RNA and is implicated in cell mobility (Figure 2C). IPA regulator effects predict EGFR to be upstream of many of the proteins effected by hydroxyacetone treatments (Figure 2D).

### Hydroxyacetone Affected Mitochondria and ROS Levels

The proteomics data indicated that mitochondria were major targets of hydroxyacetone (Figure 3). A total of 19 GO Terms related to mitochondrial physiology, various mitochondrial structures, and the respiratory chain were significantly altered by hydroxyacetone (Figure 3A). In addition, IPA Canonical pathways included Complex IV Assembly, Mitochondrial Protein Degradation, Respiratory Electron Transport, and Glucose Metabolism (Figure 3B). Complex IV included many COX enzymes that were differentially expressed in our dataset and were present in both the GO and IPA data, showing good agreement between the two analyses, and (Figure 5C).

**Figure 3:**
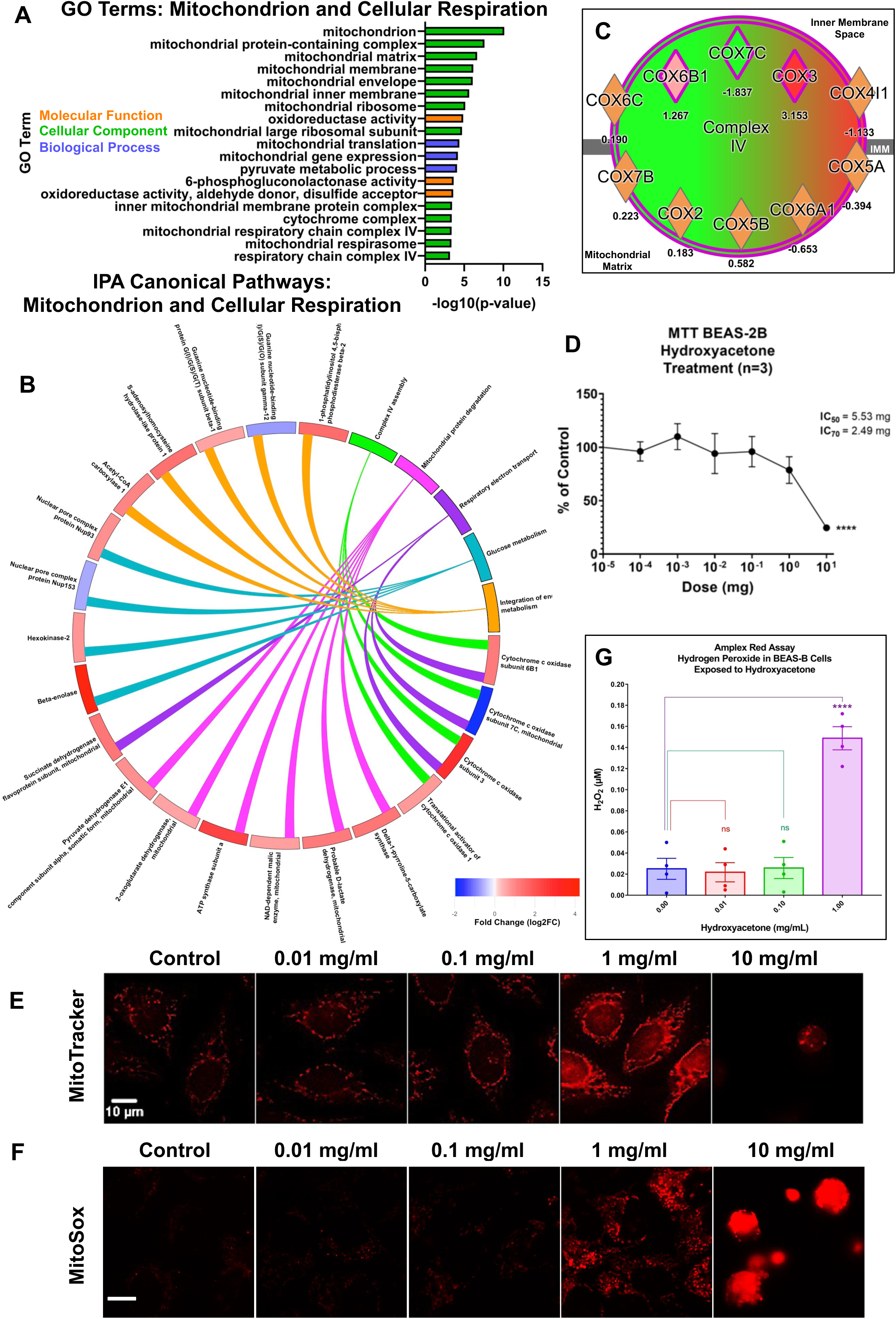
Hydroxyacetone affected mitochondrial function and generated reactive oxygen species. (A) GO Terms related to mitochondrial function and cellular respiration. (B) Circle Plot connecting mitochondrial IPA canonical pathways to their DEPs displaying fold change for each protein. (C) IPA diagram displaying proteins from Complex IV of the mitochondrial respiratory chain showing significant upregulation of COX3. (D) MTT assay for hydroxyacetone. (E) Labeling of hydroxyacetone exposed BEAS-2B cells with MitoTracker™ Red and (F) MitoSox™ Red. (G) Amplex™ Red assay detected H_2_O_2_ in the culture medium following hydroxyacetone exposure of BEAS-2B cells.

To follow-up on the proteomics data, BEAS-2B cells in monolayer cultures were treated with hydroxyacetone and examined using concentration-response assays that monitor mitochondrial endpoints or ROS production, which would be expected to increase if mitochondria are stressed (50). The cytotoxicity of hydroxyacetone was evaluated in BEAS-2B cells using the MTT assay, which measures mitochondrial reductase activity (Figure 3D). The IC_50_ and IC_70_ in the MTT assay were 5.53 and 2.49 mg/mL of hydroxyacetone, respectively. A majority of the aerosols generated from products in our sample set had concentrations of hydroxyacetone above the IC_70_.

The mitochondria were further studied with in vitro assays on BEAS-2B cells. MitoTracker™ Red is a cell-permeant dye that preferentially accumulates in mitochondria due to their negative membrane potential. Because the fluorescence of MitoTracker™ Red increases as the mitochondrial membrane potential increases, it can be used to evaluate mitochondrial activity and health. Control mitochondria had low levels of red fluorescence which did not change at the two lowest concentrations of hydroxyacetone (0.01 and 0.1 mg/mL) (Figure 3E). However, at 1.0 mg/mL, hydroxyacetone increased MitoTracker Red fluorescence significantly, indicating an increase in the mitochondrial membrane potential and hence activity (Figure 3F). At the highest hydroxyacetone concentration (10 mg/mL), red fluorescence was significantly decreased indicating weak mitochondrial membrane potential and diminished mitochondrial activity, in agreement with the MTT assay.

Because stimulation of mitochondrial activity would be expected to increase levels of superoxide, a byproduct of oxidative phosphorylation, BEAS-2B cells were treated with hydroxyacetone then labeled with MitoSox™ Red, which detects superoxide in mitochondria (Figure 3F). Micrographs showed a significant increase in MitoSox™ Red fluorescence at 1 mg/ml of hydroxyacetone, in good agreement with the MitoTracker Red data. At 10 mg/ml, cells were rounded and bright red, probably because the red fluorescence had become concentrated by cell contraction.

The MitoTracker™ Red and MitoSox™ Red labeling indicated cells were experiencing oxidative stress at 1 mg/ml of hydroxyacetone. Normally, superoxide is metabolized to hydrogen peroxide, a less toxic reactive oxygen species (ROS). When hydrogen peroxide levels become elevated, the antioxidative mechanism can become saturated and hydrogen peroxide can leak into the culture medium. To test this idea, BEAS-2B cells were incubated in various concentrations of hydroxyacetone then their culture medium was assayed for hydrogen peroxide using the Amplex™ Red assay (Figure 3G). Hydrogen peroxide was low in the control and the two low concentrations of hydroxyacetone. However, at 1 mg/ml of hydroxyacetone, the concentration that increased mitochondrial activity and superoxide production, hydrogen peroxide was significantly elevated in the culture medium.

### Hydroxyacetone Causes Depolymerization of Actin Filaments

GO and IPA analysis generate terms indicating that the actin cytoskeleton was a target of hydroxyacetone (Figure 4A) and rounding of the cells from the mitochondrial assays further supported an effect on actin (Figure 3E-F). To analyze these effects in more depth, BEAS-2B cells were exposed for 2 or 8 hours to various concentrations of hydroxyacetone (1 mg, 3.3 mg, or 10 mg/mL) then labeled using fluorescent phalloidin to visualize filamentous actin. The control had a network of long intact actin filaments in the cytoplasm, beneath the plasma membrane, and around the nucleus (Figures 4B, C). In contrast, all treatments affected BEAS-2B cell f-actin, with increased depolymerization of the filaments as hydroxyacetone concentration increased. Depolymerization was clearest and occurred first for cytoplasmic filaments and those surrounding the nucleus (Figures 4B and 4C - 1 mg/mL concentration).

**Figure 4:**
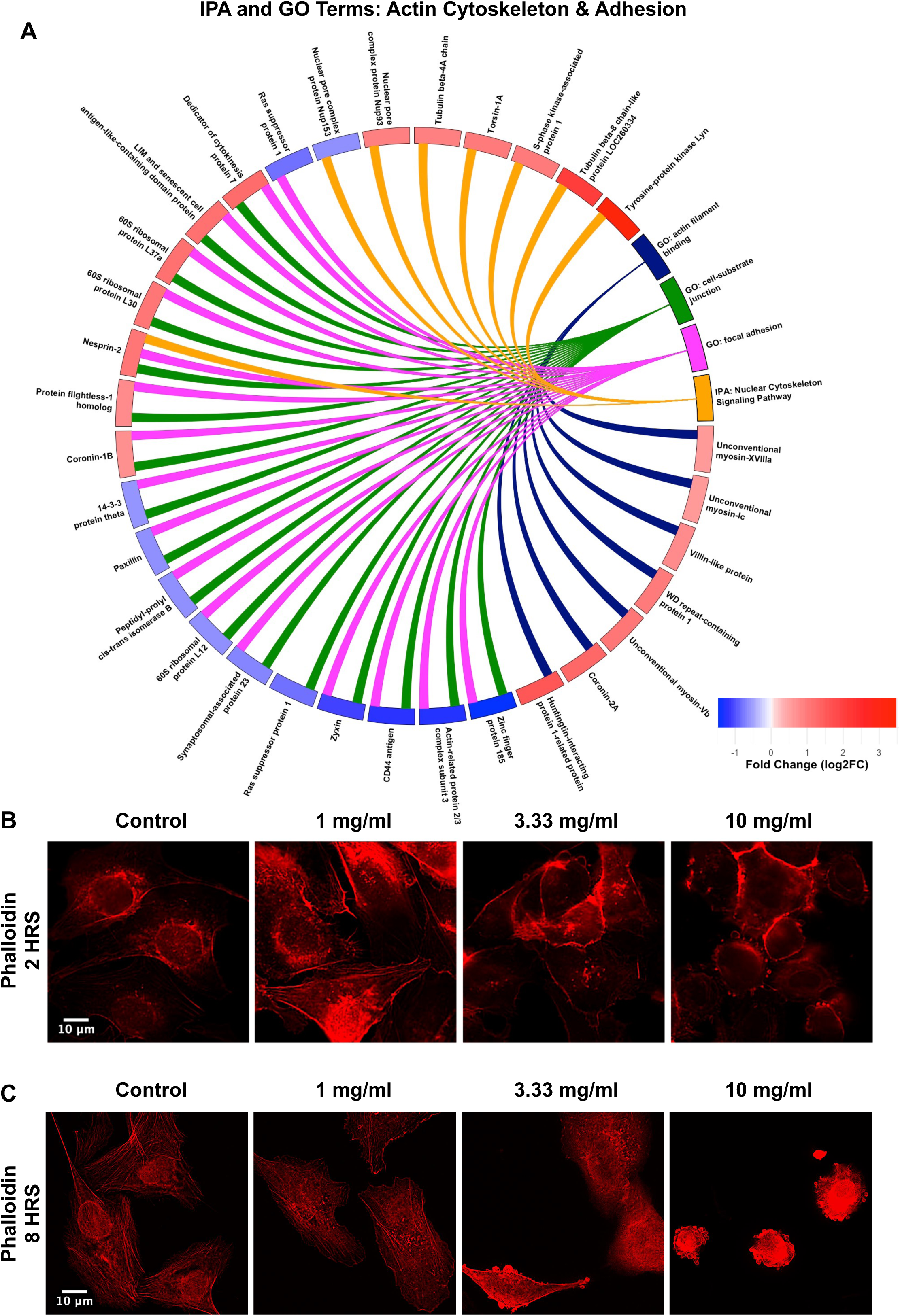
Hydroxyacetone caused depolymerization of the actin cytoskeleton. (A) Circle Plot with IPA Canonical Pathways and GO Terms related to Actin Cytoskeleton and Adhesion. Pathways and terms connect to DEPS which are color coded to show up (red) and down (blue) fold change. Phalloidin staining of BEAS-2B cells after 2 hours (B) and 8 hours (C) of treatment with hydroxyacetone. Red fluorescence indicates an increase in mitochondrial membrane potential.

**Figure 5:**
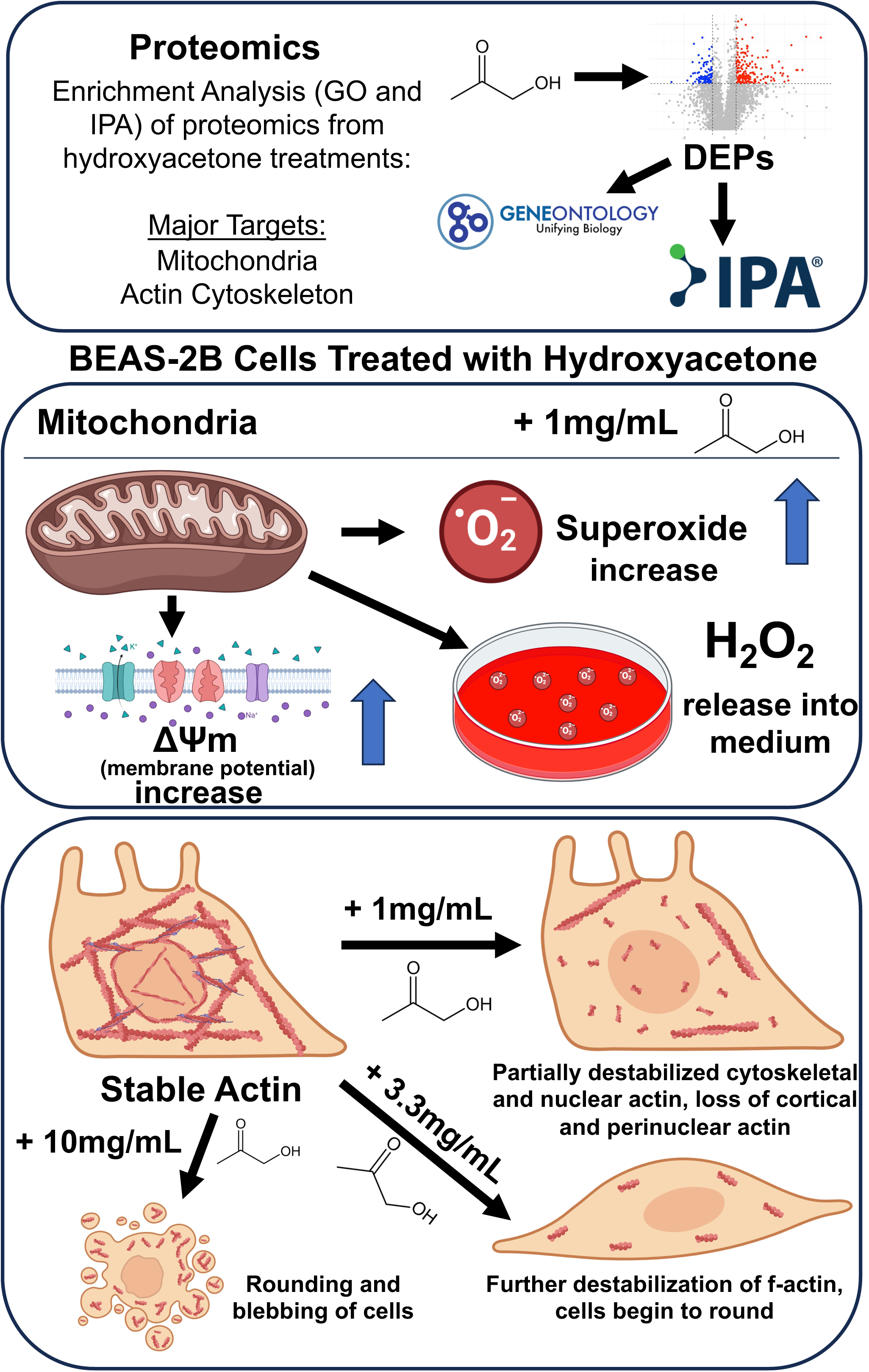
Summary of main findings. Proteomics provided a global overview of the effects produced by hydroxyacetone on EpiAirway tissue. Mitochondria and the actin cytoskeleton were two major targets of hydroxyacetone exposure identified by enrichment analysis (GO and IPA). Follow-up with BEAS-2B cells showed hydroxyacetone caused an increase in mitochondrial activity, superoxide, and hydrogen peroxide. The perinuclear actin ring and cytoplasmic actin filaments depolymerized first, while the actin filaments beneath the plasma membrane depolymerized at a higher hydroxyacetone concentration. Hydroxyacetone at 10 mg/mL caused cell blebbing and cell collapse.

Filaments beneath the plasma membrane seemed more robust to treatment and were observed at 1 mg/mL in 2-hour treatments (Figure 4B) but were gone by 8 hours of exposure (Figure 4C). At 10 mg/mL, cells rounded up and had surface blebs, which were consistent with the rounding in the MitoTracker™ and MitoSox™ data. At lower concentrations, the filaments were irregular and truncated, and cells were not as well spread as the control. These effects were observed at 2 hours but were more pronounced after 8 hours (Figure 4B-C).

## DISCUSSION

Hydroxyacetone emerged as a predominant reaction product in aerosols generated from EC fluids that users had previously associated with adverse symptoms (48). Hydroxyacetone was detected in nearly all aerosol samples and reached concentrations as high as 12Lmg/mL in aerosols from several products, exceeding the levels of all other reaction products identified in this study (48). Given its prevalence and elevated concentrations, we prioritized hydroxyacetone for toxicological characterization. Initial assessments were conducted using 3D human EpiAirway tissues, with proteomic profiling to obtain a global overview of molecular targets affected by exposure to hydroxyacetone. Two key targets—mitochondrial function and actin cytoskeletal integrity—were subsequently investigated in BEAS-2B cells using submerged culture conditions. At a concentration of 1Lmg/mL, hydroxyacetone induced cellular stress characterized by increased mitochondrial activity, elevated superoxide production, and accumulation of hydrogen peroxide in the culture medium. Concurrently, disruption of the actin cytoskeleton was observed, beginning with depolymerization at 1Lmg/mL and progressing to pronounced morphological changes, including blebbing and cell rounding, at 10Lmg/mL.

Previous studies have characterized hydroxyacetone as a reaction product formed during aerosol generation in EC devices (48,51,52). In addition to its formation in the EC aerosol phase, hydroxyacetone has been detected in the refill fluids of EC products aged 5–10 years post-use (53) and in reference solutions stored at ambient temperature in light (54), suggesting that it can also accumulate in the liquid phase over time. Thermal degradation of propylene glycol has been proposed as a plausible pathway for hydroxyacetone generation under typical EC operating conditions (37,51). Behar et al. (32) reported that aerosols generated from an Innokin iTaste MVP 3.0 device at 3V did not contain detectable hydroxyacetone, whereas operation at 5V resulted in its formation across all tested lab-made refill fluids, with concentrations ranging from approximately 0.1 mg/mL to 10mg/mL. Among these, aerosolized triacetin (2.8 mg/mL in 80% propylene glycol) yielded the highest levels of hydroxyacetone (∼10 mg/mL), underscoring the influence of both device power settings and specific flavoring agents on hydroxyacetone production. Additional flavor chemicals found to elevate hydroxyacetone levels upon aerosolization included cinnamaldehyde (155 mg/mL), menthol (84 mg/mL), and benzyl alcohol (39 mg/mL) (32). Complementary findings from studies on other carbonyl compounds suggest that flavoring agents may modulate solvent degradation pathways, thereby favoring the formation of specific carbonyl species in EC aerosols (55). Collectively, prior reports and our current findings indicate that hydroxyacetone formation is influenced by both device parameters and fluid composition.

Our proteomics analysis of EpiAirway™ tissues exposed to hydroxyacetone using ALI conditions revealed mitochondria as primary targets. The highest-scoring IPA canonical pathway was “Mitochondrial Protein Degradation,” and all significant IPA Tox List Terms were associated with mitochondrial function and oxidative stress, collectively indicating that heightened mitochondrial activity accompanied by oxidative stress can be deleterious to epithelial integrity if unresolved. A notable point of concordance between the IPA and GO enrichment analyses was the identification of Mitochondrial Respiratory Chain Complex IV as a significantly affected target. This pathway emerged as a canonical hit in IPA and was supported by two GO terms within the cellular component ontology. In our ALI proteomics dataset, three cytochrome c oxidase (COX) subunits associated with Complex IV were differentially expressed relative to the PBS control (Figure 3C). The directionality of the fold change varied among subunits, consistent with their distinct functional roles and regulatory origins. COX3 is encoded by the mitochondrial genome (MT-CO3), whereas COX6B1 and COX7C are nuclear-encoded (56), suggesting hydroxyacetone may affect both mitochondrial and nuclear gene expression. The differential regulation of COX subunits—particularly the robust upregulation of COX3—provides additional compelling evidence that hydroxyacetone exposure disrupts mitochondrial homeostasis and contributes to respiratory chain dysfunction.

Findings from the submerged monolayer cultures of BEAS-2B cells corroborated the proteomic data and demonstrated a clear concentration-dependent response of mitochondria to hydroxyacetone. Notably, 1Lmg/mL of hydroxyacetone induced a significant increase in mitochondrial membrane potential (MitoTracker™) consistent with increased ATP synthesis, elevated mitochondrial superoxide production (MitoSox™), and increased extracellular hydrogen peroxide levels (Amplex Red). These converging endpoints suggest that 1 mg/mL of hydroxyacetone exposure stressed cells leading to increased oxidative phosphorylation accompanied by a robust oxidative stress response. In contrast, the MTT assay did not detect a significant effect at 1Lmg/mL, likely due to its lower sensitivity compared to the fluorescence-based mitochondrial assays. Our observation aligns with a previous report that also found no significant MTT response to hydroxyacetone at 1Lmg/mL in human pulmonary fibroblasts (hPF) and A549 cells (32).

The cytoskeleton was another major target of hydroxyacetone in the proteomics dataset based on GO terms, such as “actin filament binding”, and IPA canonical pathway analysis. Notably, “Nuclear Cytoskeleton Signaling” emerged as the second highest-scoring IPA canonical pathway, suggesting that hydroxyacetone may disrupt cytoskeletal architecture at both the cytoplasmic and nuclear levels. This idea was supported by in vitro experiments with BEAS-2B cells, which demonstrated concentration-dependent actin filament depolymerization following hydroxyacetone exposure. Cytoplasmic actin filaments and the perinuclear actin ring were visibly disrupted at 1Lmg/mL as early as 2 hours post-treatment. In contrast, cortical actin filaments were more resistant, exhibiting only partial depolymerization at 3.3Lmg/mL. Complete loss of filamentous actin and cellular collapse, followed by apparent blebbing were observed after 8 hours of exposure to 10Lmg/mL of hydroxyacetone.

These findings reveal differential sensitivities among actin filament populations and are consistent with cytoskeletal stress induced by hydroxyacetone. Importantly, depolymerization of the perinuclear actin ring may compromise nuclear shape and disrupt mechanical coupling between the nucleus and cytoskeleton. Supporting this, several proteins involved in nuclear– cytoskeletal interactions were differentially expressed in the proteomics dataset, including nuclear pore complex components (Nup153 and Nup93) and Nesprin-2, which links the nuclear envelope to the perinuclear ring actin (57). Comparable cytoskeletal disruption has been reported for other EC aerosol constituents. For example, the synthetic coolant WS-23 induced actin filament depolymerization in BEAS-2B cells, with perinuclear ring disruption observed at concentrations as low as 0.045Lmg/mL (26). Although not part of our BEAS-2B study, the proteomics data also indicated that tubulin proteins (e.g., tubulin βA chain) may likewise have been perturbed by hydroxyacetone.

Very limited data exist on the inhalation toxicity of hydroxyacetone, while structurally related compounds, such as dihydroxyacetone, have been more extensively studied. Although the U.S. Food and Drug Administration approved dihydroxyacetone for topical use in cosmetic products such as self-tanning formulations in 1977, it explicitly cautions against inhalation or exposure to the eyes and nasal mucosa due to its irritant properties (58,59). Dihydroxyacetone has been detected in EC aerosols (60), albeit at substantially lower concentrations than hydroxyacetone in our sample set (48). Inhalation exposure to dihydroxyacetone has been associated with mucosal irritation and genotoxicity (58,61–65).

Experimental studies have demonstrated that dihydroxyacetone exerts toxic effects on human airway epithelial cells cultured at the ALI, including inhibition of ciliary beating, reduced MUC5AC secretion, and diminished MMP9 release (63). Additional reports describe dihydroxyacetone-induced cell cycle arrest and apoptosis in melanoma cells (66), genotoxicity and chromosomal aberrations in BEAS-2B cells (64), and pulmonary injury and fibrosis in murine models (65). Notably, dihydroxyacetone also induced mitochondrial stress and autophagy in HEK293T cells (62), findings that parallel our observations with hydroxyacetone in bronchial epithelial models.

The concentrations of hydroxyacetone that elicit adverse symptoms in humans remain poorly defined, complicating direct extrapolation of our findings to EC exposure scenarios. According to publicly available health and safety documentation (67), inhalation of hydroxyacetone vapors may induce symptoms such as drowsiness, dizziness, and respiratory irritation. At unspecified high concentrations, reported effects include pulmonary irritation with coughing and nausea, central nervous system depression manifesting as headache and dizziness, slowed reflexes, fatigue, and incoordination. Despite these accounts, hydroxyacetone is not formally classified as “harmful by inhalation,” likely due to insufficient animal and human toxicological data. Notably, several symptoms described in these safety sheets—nausea, dizziness, cough, headache, shortness of breath, and fatigue—closely mirror those reported by users of the EC products analyzed in our study (48). This overlap raises the possibility that some symptomatic EC users may have experienced adverse effects resulting from inhalation of elevated hydroxyacetone concentrations.

## Conclusion

Hydroxyacetone is a prevalent reaction product in EC aerosols, with concentrations reaching 12 mg/mL across diverse formulations. Despite its widespread occurrence and frequent high abundance, hydroxyacetone has received limited attention in the toxicological literature. Proteomic analysis of 3D EpiAirway tissue exposed at the ALI revealed broad differential expression of proteins, with mitochondrial pathways and the actin cytoskeleton identified as primary targets. These findings were corroborated by in vitro toxicological assays, which demonstrated that hydroxyacetone induced mitochondrial dysfunction, oxidative stress, and actin destabilization in human bronchial epithelial cells in a concentration-dependent manner. Its occurrence at elevated concentrations in select EC products may have contributed to the adverse health effects reported by users. Collectively, these results highlight hydroxyacetone as a biologically active aerosol constituent and underscore the need for further investigation into its inhalation toxicology and potential contribution to EC-related respiratory effects.

## Limitations

Hydroxyacetone concentrations in EC aerosols are highly sensitive to variations in device temperature, power output, and user topography, which likely contribute to the wide range of concentrations reported across studies, including those observed with the products used in our experiments. In the present study, exposures were acute in both the EpiAirway tissue model and BEAS-2B cell assays. It is plausible that chronic exposure may yield distinct biological outcomes, and that hydroxyacetone could exert toxicological effects at lower concentrations under prolonged exposure conditions. In studies dealing with single chemical exposures, there is always the possibility that the treatment chemical is metabolized by the exposed tissue to another chemical(s), which produces the observed effect.

## Author Contributions

P.T. and M.W. formed the concept and design of this study. M.W., T.M. M.H and N.H. carried out sample preparation, data collection, processing, and analysis. M.W. and P.T wrote the first draft of the manuscript. T.M., M. H., M.W. and P. T. edited the manuscript.

## Notes

The authors declare no competing financial interest.

## Acknowledgements

We would like to thank Dr. James F. Pankow, Dr. Wentai Liu, and Kevin McWhirter for analyzing hydroxyacetone concentrations deposited in the VitroCell™ Cloud Chamber wells during ALI exposure.

---

Supplemental Table 1: Table containing products reported to cause symptoms in users. Hydroxyacetone was detected in aerosols at much higher concentrations than in refill fluids.

Supplemental Table 2: Proteomics data used for enrichment analysis.

Supplemental Figure 1: (A) Concentrations of hydroxyacetone quantified in VitroCell™ Cloud Chamber wells, these were the concentrations of hydroxyacetone that the individual inserts received in a single exposure. (B) Top differentially expressed proteins from hydroxyacetone treatments.

Supplemental Figure 2: Circle plot connecting top IPA Canonical Pathways (z > 2) connected to DEPs in each pathway. Fold change of proteins is displayed by the protein name (red = upregulated, blue = downregulated) and –log(p-value) is reflected based on size of the dot by the pathway.

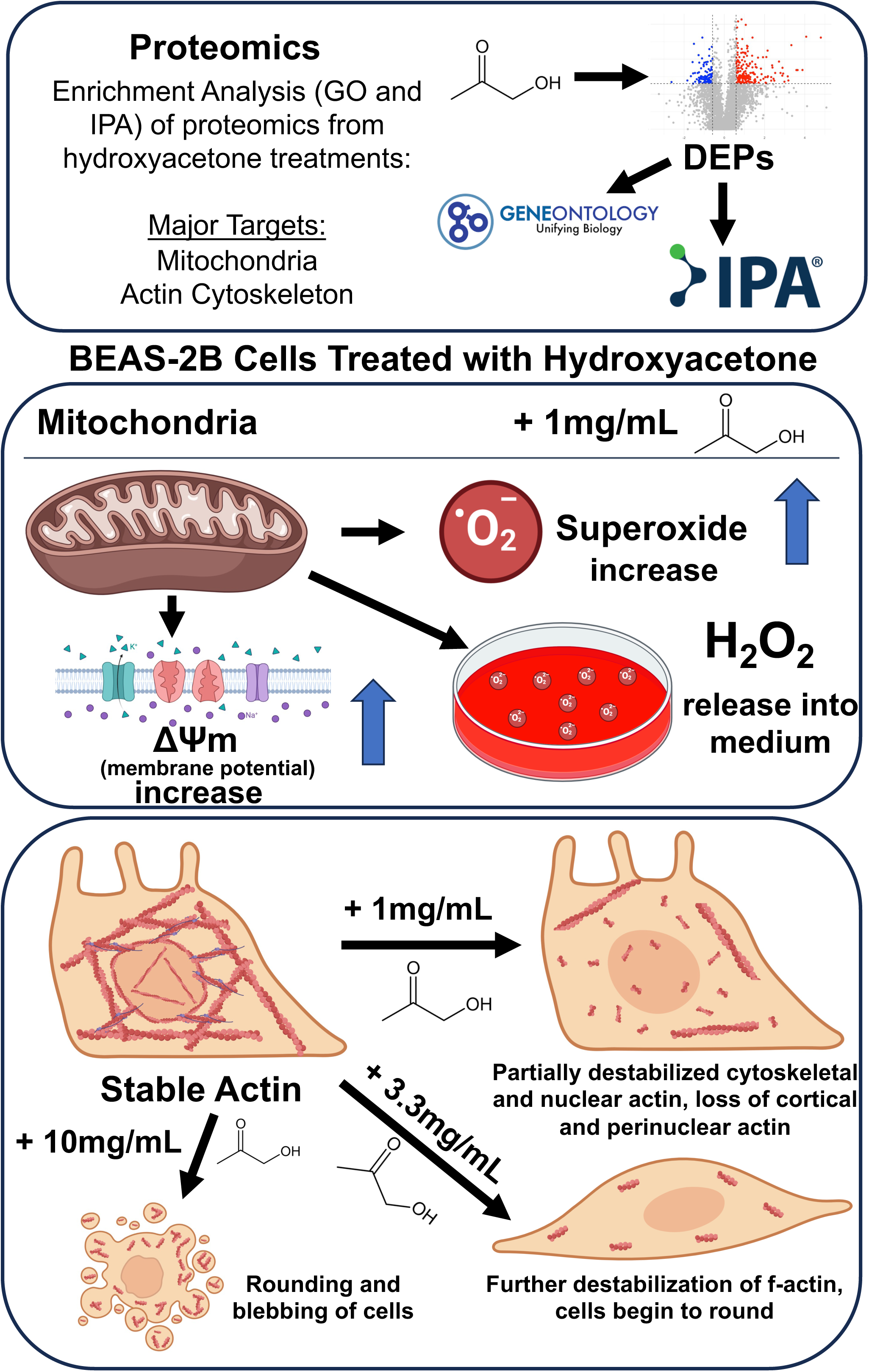

