## Supplemental Figures 1 and 2 for "Unraveling the Toxicological Effects of Hydroxyacetone - A Reaction Product in Electronic Cigarette Aerosols"

**A**

| | Well 1 (ug/mL) | Well 2 (ug/mL) | Well 3 (ug/mL) | Well 4 (ug/mL) | Well 5 (ug/mL) | Mean ± SD (ug/mL) |
|---|---|---|---|---|---|---|
| **Hydroxyacetone** | **106.7** | **120.3** | **116.1** | **102.8** | **111.6** | **111.5±6.3** |

**B**

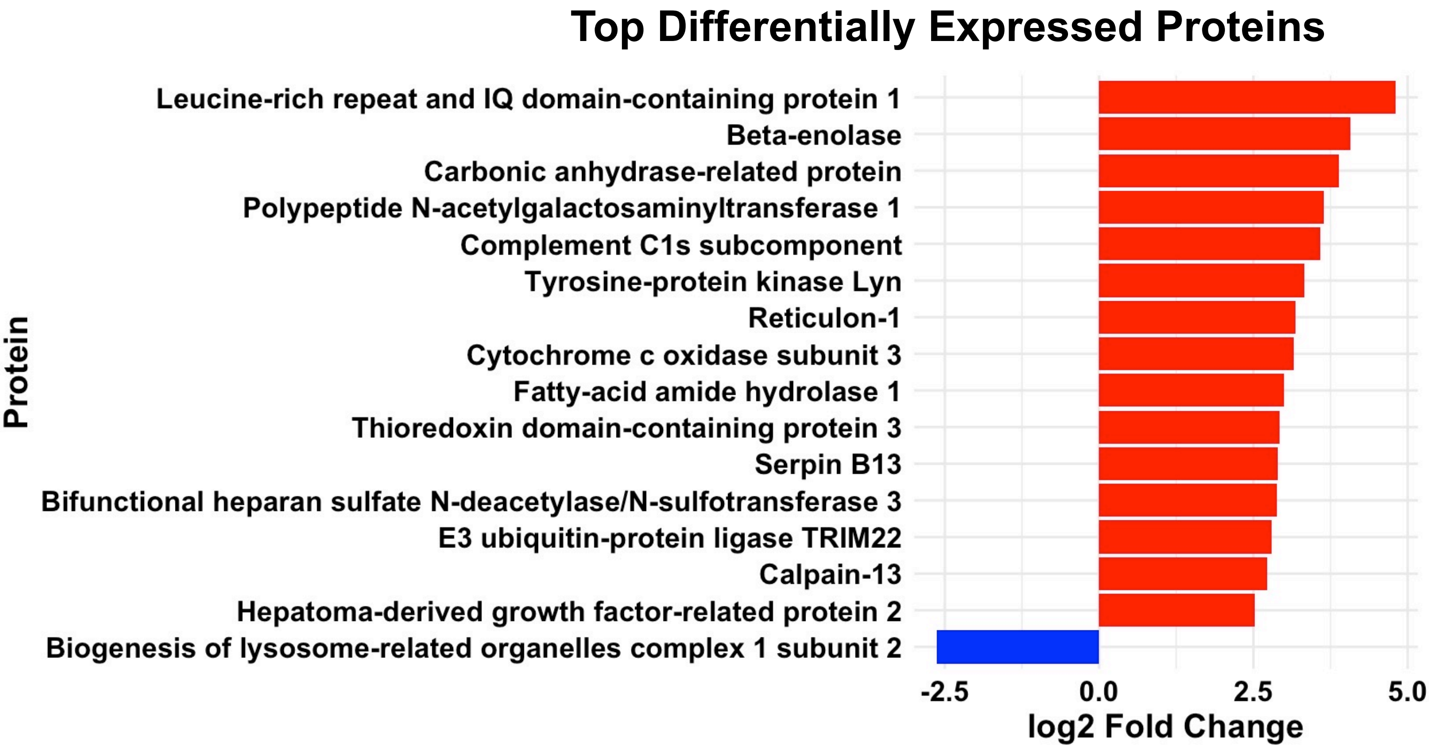

**Top Canonical Pathways (z < 2)**

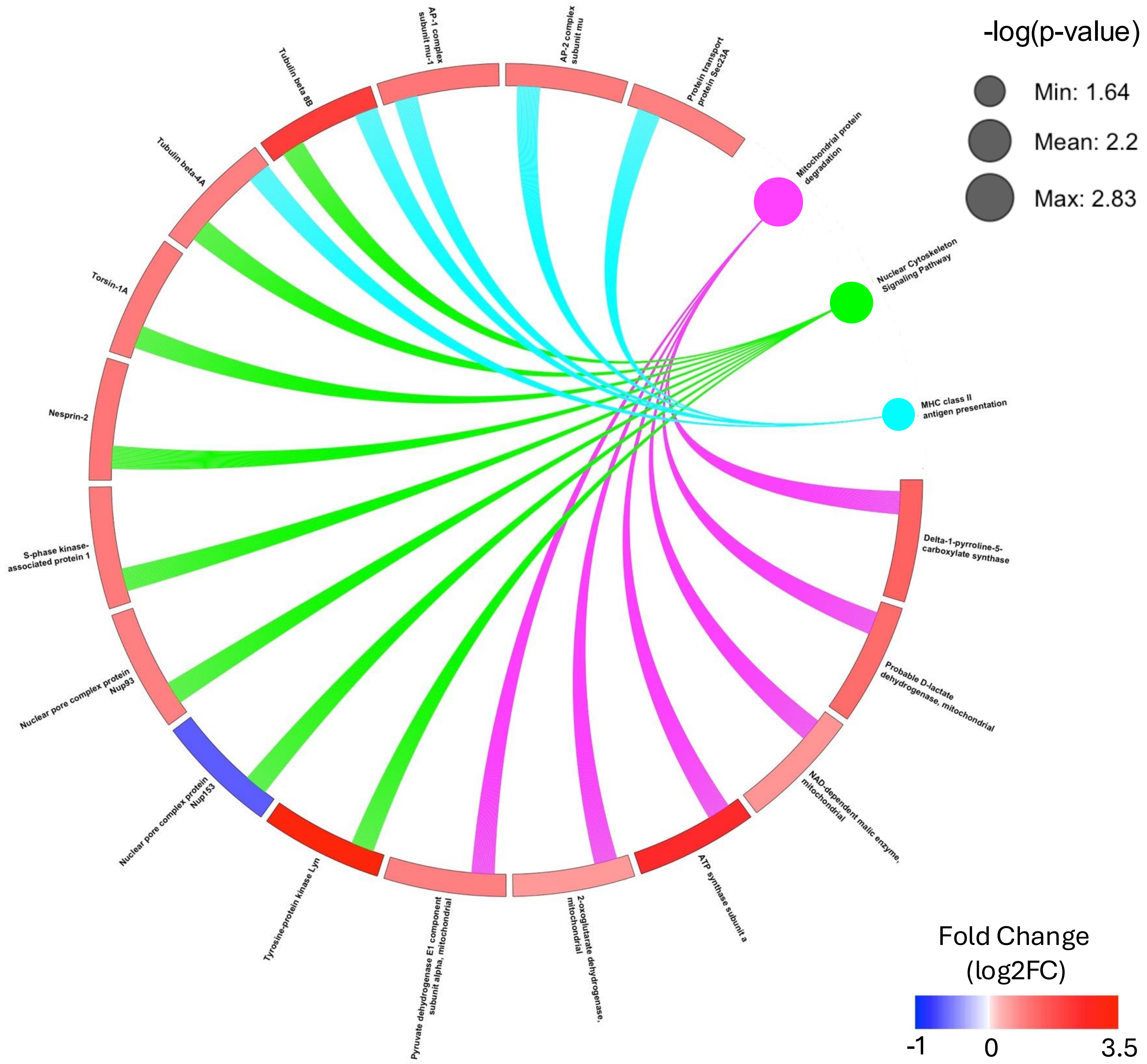
